# SCYN: Single cell CNV profiling method using dynamic programming

**DOI:** 10.1101/2020.03.27.011353

**Authors:** Xikang Feng, Lingxi Chen, Yuhao Qing, Ruikang Li, Chaohui Li, Shuai Cheng Li

## Abstract

Copy number variation is crucial in deciphering the mechanism and cure of complex disorders and cancers. The recent advancement of scDNA sequencing technology sheds light upon addressing intratumor heterogeneity, detecting rare subclones, and reconstructing tumor evolution lineages at single-cell resolution. Nevertheless, the current circular binary segmentation based approach proves to fail to efficiently and effectively identify copy number shifts on some exceptional trails. Here, we propose SCYN, a CNV segmentation method powered with dynamic programming. SCYN resolves the precise segmentation on two *in silico* datasets. Then we verified SCYN manifested accurate copy number inferring on triple negative breast cancer scDNA data, with array comparative genomic hybridization results of purified bulk samples as ground truth validation. We tested SCYN on two datasets of the newly emerged 10x Genomics CNV solution. SCYN successfully recognizes gastric cancer cells from 1% and 10% spike-ins 10x datasets. Moreover, SCYN is about 150 times faster than state of the art tool when dealing with the datasets of approximately 2000 cells. SCYN robustly and efficiently detects segmentations and infers copy number profiles on single cell DNA sequencing data. It serves to reveal the tumor intra-heterogeneity. The source code of SCYN can be accessed in https://github.com/xikanfeng2/SCYN. The visualization tools are hosted on https://sc.deepomics.org/.

## Background

Numerous studies have shown that copy number variations(CNV) can cause common complex disorders (1–5). Copy number aberration (CNA), aka, somatic CNV, is also reported to be a driving force for tumor progression and metastasis. For example, George *et al* reported the high amplification of oncogene gene *PD-L1* in small-cell lung cancer (6) and amplification of *MYC* is announced prevailing in pan-cancer studies (7). The loss of tumor suppressor genes like *KDM6A* and *KAT6B* are proclaimed indirectly amplifies harmful cancer-related pathways (8, 9).

Conventional experimental protocols for CNV segmentation lies in the following scenarios. Researchers may infer a coarse CNV profiles utilizing bulk RNA sequencing (10) and single cell RNA sequencing (11–13). Moreover, scientists may leverage bulk genome such as DNA array comparative genomic hybridization (aCGH) (14), single-nucleotide polymorphism (SNP) arrays (15, 16), and DNA next generation sequencing (NGS) (17, 18) to generate high resolution CNV. Although bulk genome sequencing studies have contributed insights into tumor biology, the data they provide may mask a degree of heterogeneity (19). For instance, if the averaged read-out overrepresents the genomic data from the dominant group of the tumor cells, rare clones will be masked from the signals. The advent of single-cell DNA (scDNA) sequencing delivers a potential solution (20–22). Researchers can overwhelm the deficiencies of bulk approaches to address intratumor heterogeneity (ITH) (22), detect rare subclones (19), and reconstruct tumor evolution lineages (20, 23).

In this study, we concentrate on the CNV segmentation and turning points detection approaches customized for single cell DNA sequencing. CNV Segmentation refers to partitioning the genome into non-overlapping segments with the objective of that each segment shares intra-homogeneous CNV profile, and the segment boundaries are often termed to be checkpoints or turning points (24). Although numerous CNV segmentation tools have emerged leveraging high throughput sequencing data such as Circular Binary Segmentation (CBS) (25, 26) and Hidden Markov Model (HMM) (27, 28), the methods customized for scDNA data is in its infancy. Gingko (29), SCNV (30), and SCOPE (31) applied diverse strategies to normalize the scDNA intensities through simultaneously considering sparsity, noise, and cell heterogeneity, and adopted variational CBS for checkpoint detection. While after *in silico* experiments, we argue that those CBS approaches might not lead to an optimal segmentation result, some turning points might be masked. Furthermore, with the advance of large scale high throughput technologies, the scale of cells for a single dataset climbs exponentially. For instance, the newly emerged 10x Genomics CNV solution can profile the whole genome sequencing of thousands of cells at one time (22). Thus, efficiently processing scDNA-seq data is crucial. However, current scDNA CNV segmentation methods are too time-consuming to process thousands of cells.

Therefore, in this paper, we propose SCYN, an effecient and effective dynamic programming approach for single cell data CNV segmentation and checkpoint detection. SCYN resolves the precise turning points on two *in silico* datasets, while existing tools fail. SCYN manifested more precise copy number inference on a triple-negative breast cancer scDNA dataset, with array comparative genomic hybridization results of purified bulk samples as ground truth validation. We tested SCYN on two datasets of the newly emerged 10x Genomics CNV solution. SCYN successfully recognizes gastric cancer cells from 1% and 10% spike-ins 10x datasets. Last but not least, SCYN is about 150 times faster than state of the art tool when dealing with thousands of cells.

## Results

### Overview of SCYN

We developed an algorithm, SCYN, that adopts a dynamic programming approach to find optimal single-cell CNV profiles. The framework for SCYN displayed in **Figure 1A**. First, the raw scDNA-seq reads of FASTQ format are pre-processed with standard procedures (see **Figure 1A**). SCYN then takes the aligned BAM files as the input. SCYN integrates SCOPE (31), which partitions chromosomes into consecutive bins and computes the cell-by-bin read depth matrix, to process the input BAM files and get the raw and normalized read depth matrices. The segmentation detection algorithm is then performed on the raw and normalized read depth matrices using our dynamic programming to identify the optimal segmentation along each chromosome. The segmentation results are further applied to copy number calculation. Finally, SCYN outputs the cell-by-bin copy number matrix and the segmentation results of all chromosomes for further CNV analysis.

**Fig. 1.**
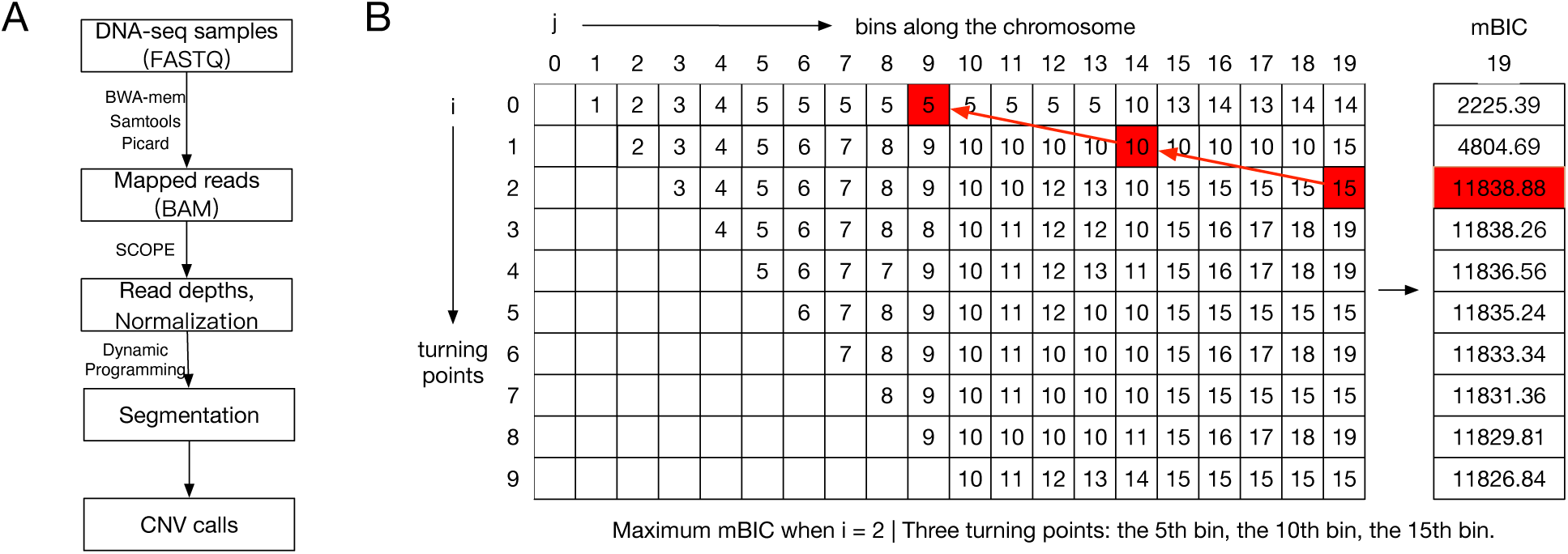
Overview of SCYN.

### SCYN effectively identifies all breakpoints on synthetic trials

To evaluate the segmentation power of SYCN against SCOPE, we generated two different combinations of the CNV intensities of blue cell and orange cell along 200 bin regions. In the first simulation, the ground truth segmentation are (1, …, 49), (50, …, 99) (100,…, 149), (150,…, 200); and the copy number state alternates between haploid and diploid. Figure 2 shows the SCOPE unable to detect the turning point 100 here, leading to erroneously dropping the loss of heterogeneity event of bin range [100, 123]. In contrast with SCOPE, SCYN accurately detected all turning points and assigned the correct copy number to all bin regions. Then, with fixed copy number turning points (50, 100, and 150) and copy number state alternates between one and four, we simulated the situation where blue cell and orange cell are always heterogeneous. In Figure 2, SCYN successfully categorized all turning points and copy number states with 100% accuracy and uncovered the cell heterogeneity. Even though SCOPE assigned correct copy number to each bin region, we found that it output five turning points 50, 100, 143, 146, and 150. In other words, SCOPE considered there exited consecutive copy number shifts among bin ranges [101, 143], [144, 146], and [147, 150], which opposite against the homogeneous fact. As previously mentioned, the core principle of CNV segmentation is partitioning the genome into non-overlapping areas with the objective of that each area shares intra-homogeneous CNV profile (24, 30). SCOPE fails to hit the correct answer as its turning point detection fails. Overall, these two experiments on synthetic data suggest that empowered with dynamic programming, SCYN can achieve the correct copy number turning point detection against the segmentation schema SCOPE proposed.

**Fig. 2.**
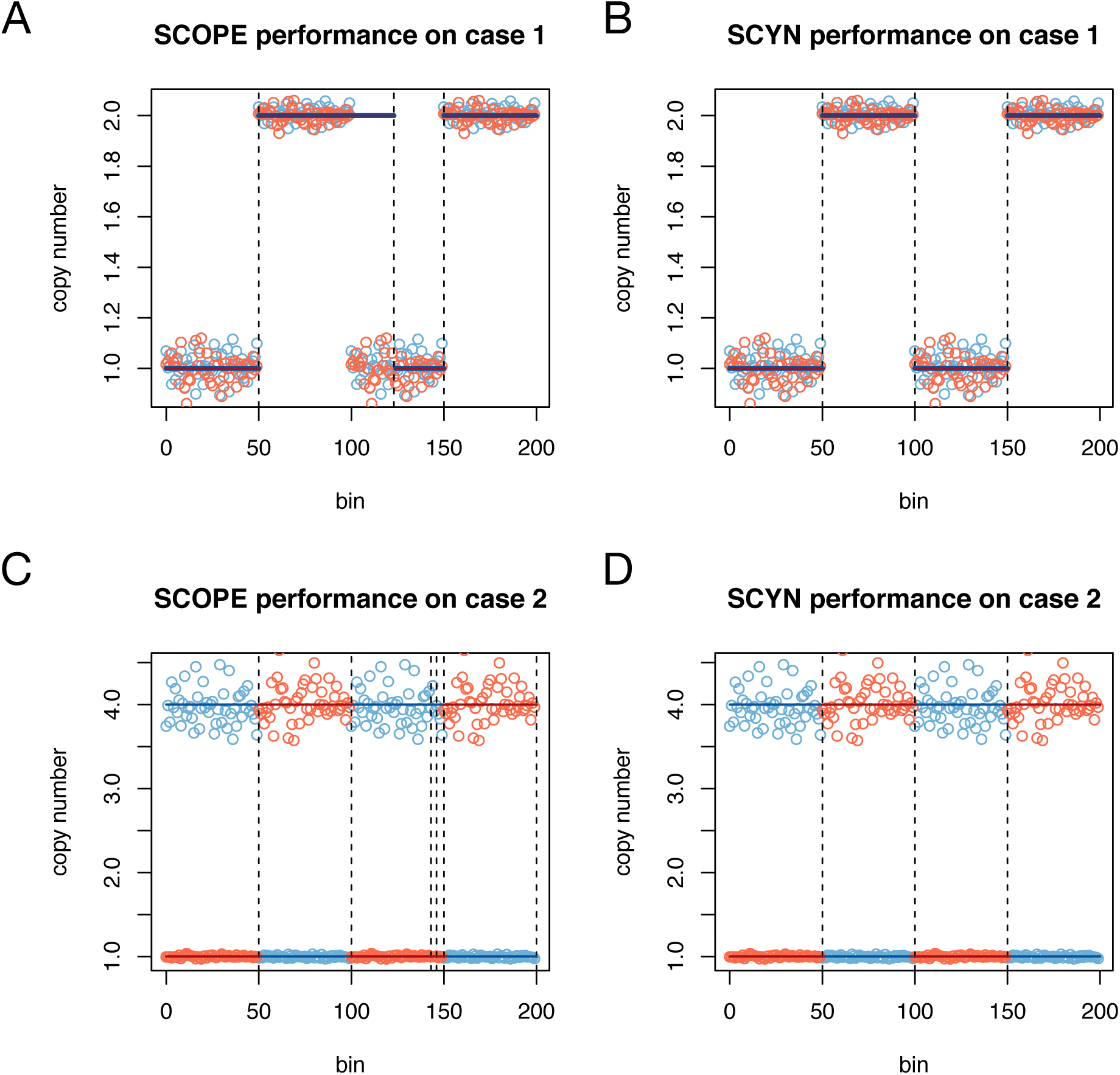
SCYN performance on synthetic cases. Hollow circles and horizontal lines denote the copy number before and after smoothing respectively. Vertical dashed lines signify the detected turning points. Orange and blue refer to cell 1 and cell 2 respectively.

### SCYN successfully identifies subclones in wet-lab cancer datasets

We illustrate the performance of SCYN in cancer single-cell datasets. We collected two cancer data sets, namely the Nature_TNBC (two triple-negative breast cancers) (32) and 10x_Gastric (gastric cancer spike-ins). We illustrated the tumor intra-heterogeneity discovered by SCYN and validated the results of SCYN against the estimation made by SCOPE for ground truth available datasets.

The first benchmark dataset we investigated is Nature_TNBC. 100 single cells were separately sequenced from two triple-negative breast cancer samples, namely, T10 and T16 (32). For T10, we removed cell SRR054599 as it did not pass the quantity control, resulting 99 single cells from held four subgroups: Diploid (D), Hypodiploid (H), Aneuploid A (A1), and Aneuploid B (A2). We first verified if SYCN could replicate the subclone findings previously reported. Figure 3A demonstrates the genome-wide copy number profiles across the 100 single cells for T10. Overall, the cell subclones recognized by SCYN are concordant with the outputs of SCOPE (see Additional file 1, Supplementary Figure S1A) and Navin *et al*.’s findings. With hierarchical clustering, SCYN categorizes T10 into seven clusters. As illustrated in Figure3 and Additional file 1 Supplementary Figure S2A-3A, for T10, cluster 1 matches the diploid (D) cells and cluster 3 represents the hypodiploid (H) group. There are two hyperdiploid subgroups. Cluster 4 corresponds to aneuploid A (A1) and cluster 2,5,6,7 together represents aneuploid B (A2). Navin *et al*. also separately profiled the four subgroups through array comparative genomic hybridization (aCGH) (33), here we regarded the CNV profiled from aCGH as golden-standard to examine the SYCN and SCOPE performance. As illustrated in Figure 3B-C, SCYN owns a higher Pearson correlation and a lower root mean squared error (RMSE) of ground-truth against SCOPE.

**Fig. 3.**
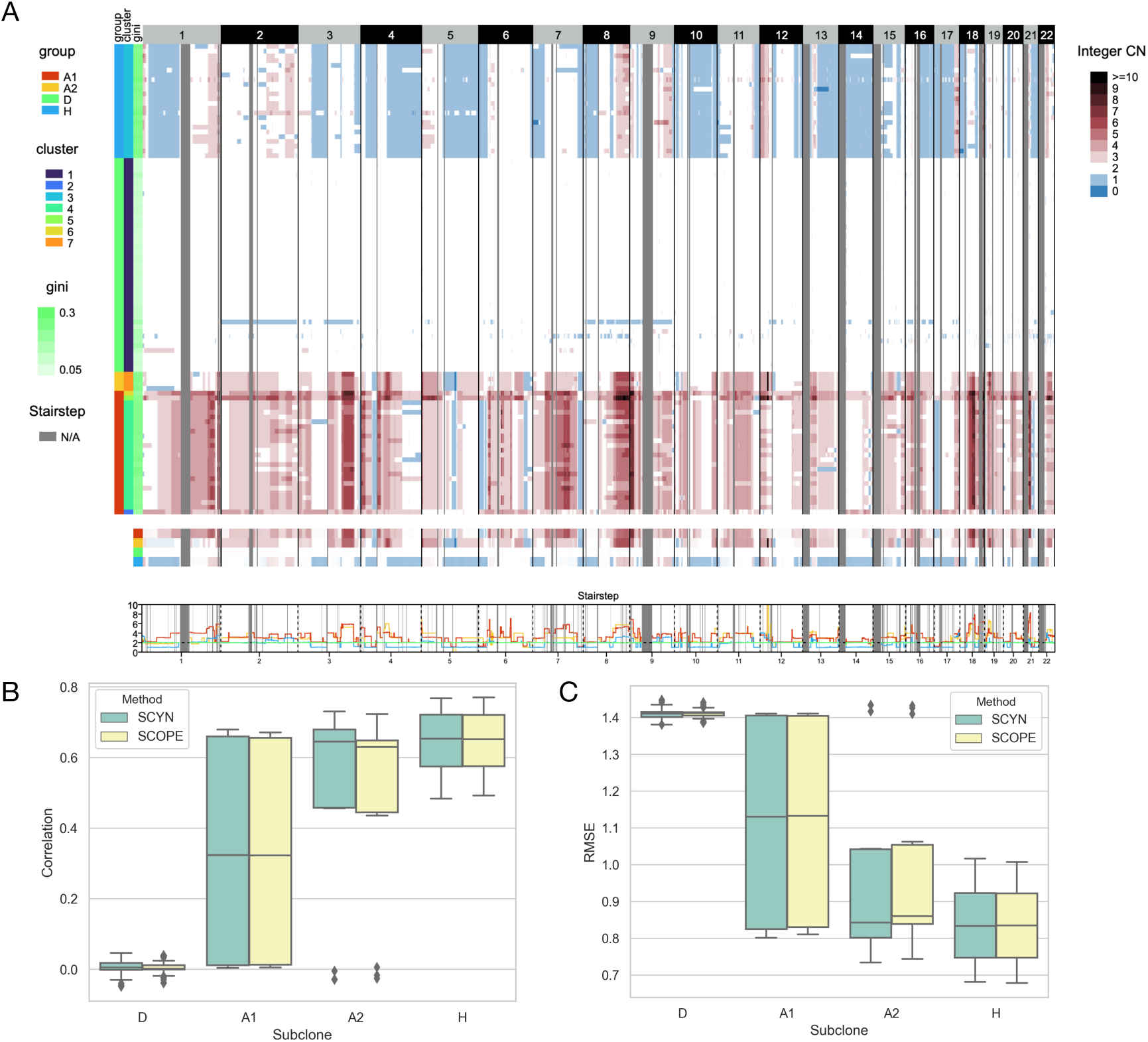
Performance of SCYN on T10. (A) Heatmap of whole genome CNV profiles (B-C) Pearson correlation and RMSE as evaluation metrics comparing results by SCYN and SCOPE against aGCH.

T16 sample is a mixture of one primary breast tumor (T16P, 52 single cells) and its corresponded liver metastasis (T16M, 48 single cells). Navin *et al*. identified five cell sub-populations: Primary Diploid (PD), Primary Pseudodiploid (PPD), Primary Aneuploid (PA), Metastasis Diploid (MD), and Metastasis Aneuploid (MA). Figure 4A records T16 genome-wide copy number profiles across the 100 single cells. In all, the cell subclones recognized by SCYN are consistent with SCOPE (see Additional file 1, Supplementary Figure S1B) and Navin *et al*.’s findings. Hierarchical clustering characterizes T16 into seven subgroups. As depicted in Figure4 and Additional file 1 Supplementary Figure S2B-3B, cluster 1 mates the primary diploid (PD) cells. Cluster 3 represents metastasis aneuploid (MA), and cluster 6,7 together pictures primary aneuploid (PA). As Navin *et al*. only profiled four bulk dissections using of T16 aCGH (33), there lacks the CNV gold standard for 16T *in su* subclones. So we calculated the CNV correlation and RMSE between inferred primary aneuploid (PA) subpopulation and the four dissections, respectively. From Figure 4B-C, although the association between PA group and four bulk dissections is relatively low, SCYN profiles a closer correlation than SCOPE with higher correlation and lower discrepancy.

**Fig. 4.**
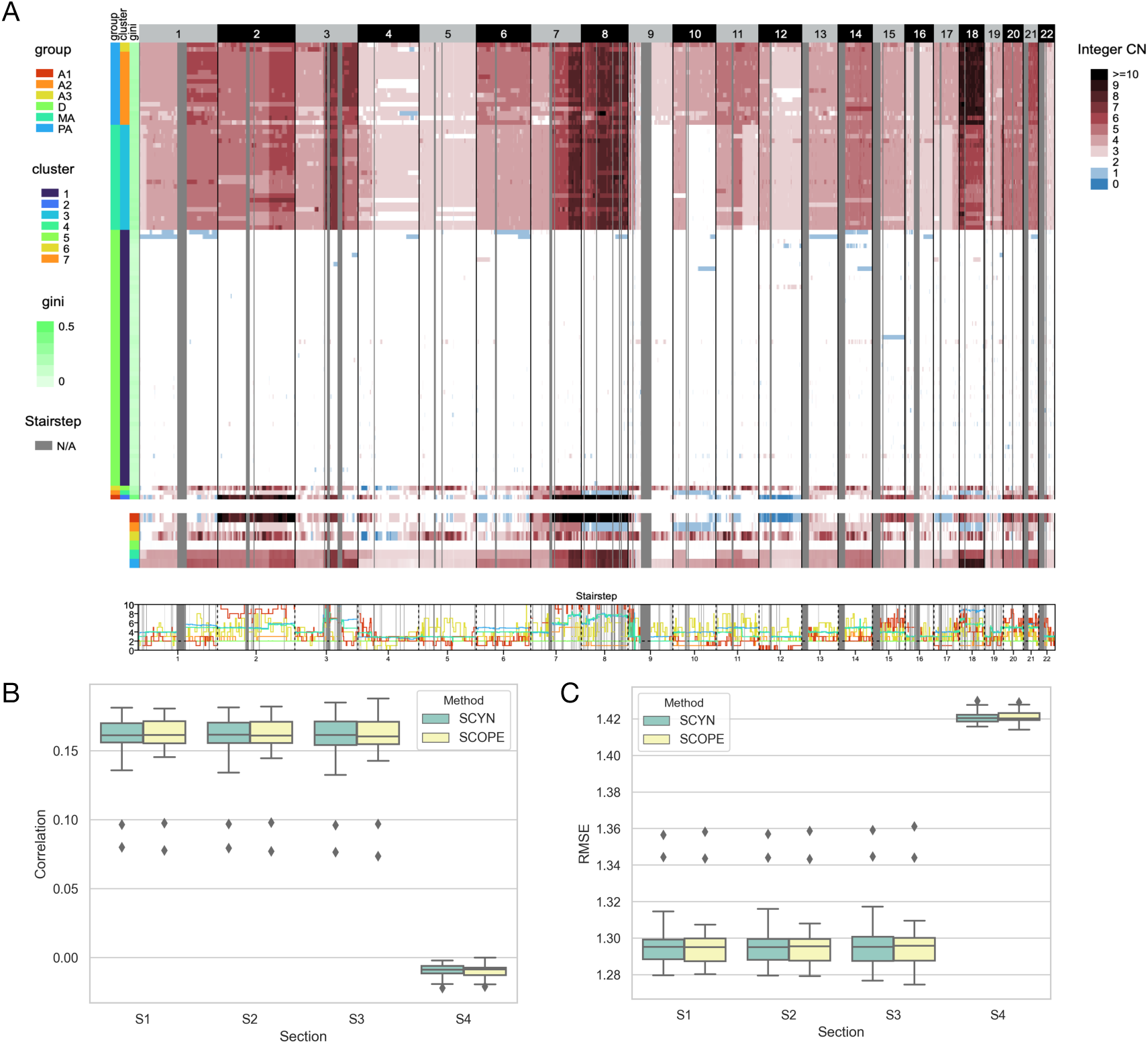
Performance of SCYN on T16. (A) Heatmap of whole genome CNV profiles (B-C) Pearson correlation and RMSE as evaluation metrics comparing results by SCYN and SCOPE against aGCH.

We next employed SCYN and SCOPE to the lately published single cell DNA spike-in demo datasets available at the 10x Genomics official website. 10x Genomics mixed BJ fibroblast euploid cell line with 1% and 10% spike-in of cells from MKN-45 gastric cancer cell line. As illustrated in the CNV heatmap Figure5A and Additional file 1 Supplementary Figure S4, SCOPE successfully distinguished the two spike-in gastric cancer cells. Furthermore, we visualized the first two principal components of the estimated CNV profiles in Figure5B-C. Cells whose Gini coefficient more massive than were highlighted in yellow and regarded as gastric cancer cells from the 1% and 10% spike-ins, respectively. Then, we checked if SYCN produced CNV profiles better preserves the cell subpopulation information against SCOPE. Leveraging Gini 0.12 as the cut-off value, we partitioned cells into normal and cancer subset as benchmark labels. Next, we practiced hierarchical clustering into CNV matrices attained from SYCN and SCOPE, and get two clusters for each spike-in sets. Then, we adopt four metrics to inquire about the clustering accuracy of SYCN against SCOPE. The adjusted Rand index (ARI) (34), Normalized mutual information (NMI) (35), and Jaccard index (JI) (36) measures the similarity between the implied groups and golden-standard labels; a value approaching 0 purports random assignment, and one reveals accurate inferring. As evidenced in Table 1 and Table 2, with ARI, NMI, and JI as measurements, SYCN holds equal clustering accuracy to SCOPE on both 1% and 10% spike-in sets, which indicates SYCN captures substantial interior tumor heterogeneity.

**Fig. 5.**
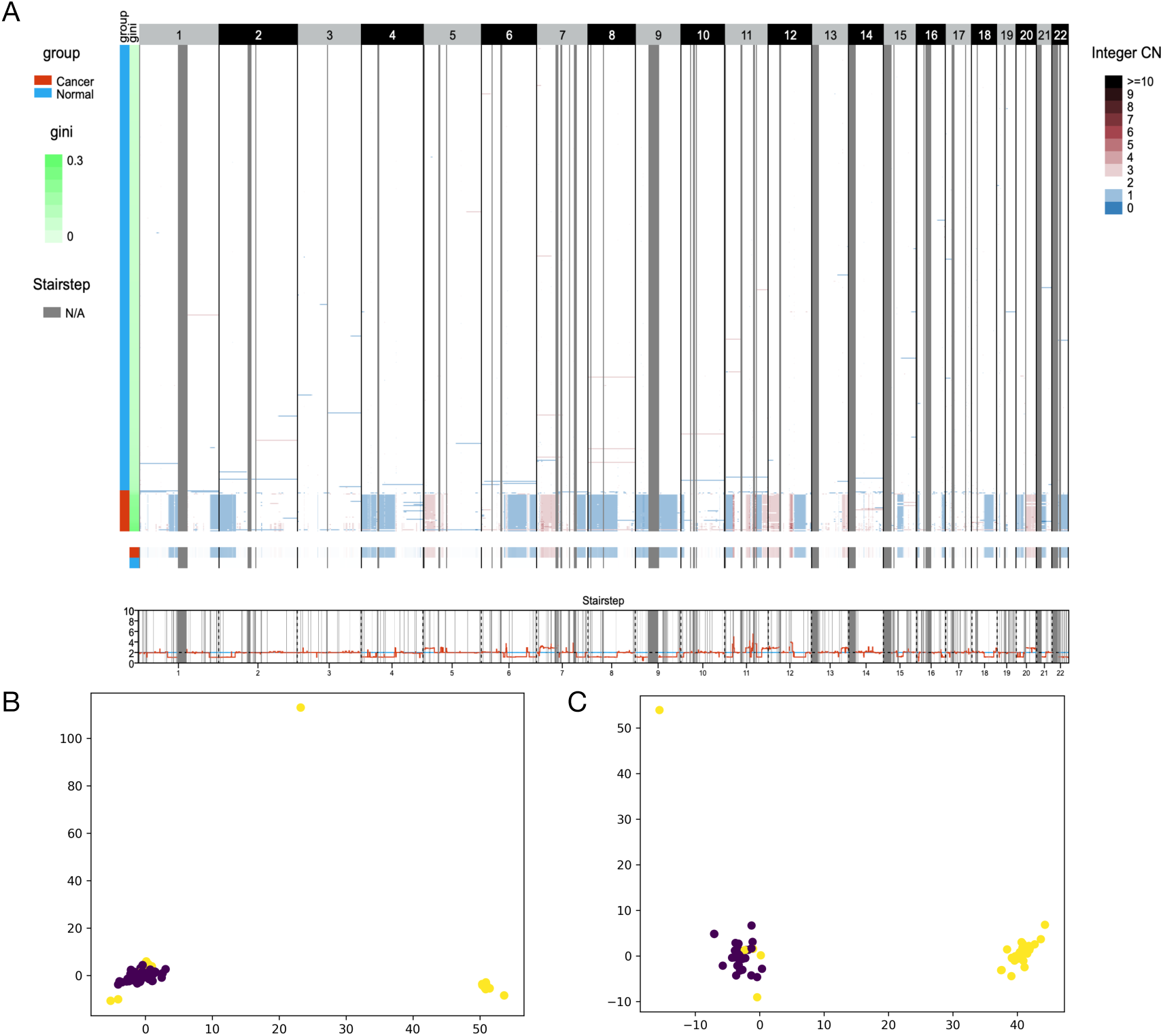
Performance of SCYN on 10x spike-ins. (A) Heatmap of whole genome CNV profiles of 10% spike-in dataset (B-C) PCA plots on 1% and 10% spike-in datasets respectively. The yellow and purple dots denote cancer and normal cell respectively.

**Table 1.** 10x 1% spike-in datasets clustering evaluation.

| Method | ARI | NMI | JI |
| --- | --- | --- | --- |
| SCYN | 0.67650 | 0.7623 | 0.5238 |
| SCOPE | 0.67650 | 0.7623 | 0.5238 |

**Table 2.** 10x 10% spike-in datasets clustering evaluation.

| Method | ARI | NMI | JI |
| --- | --- | --- | --- |
| SCYN | 0.9139 | 0.8770 | 0.8718 |
| SCOPE | 0.9139 | 0.8770 | 0.8718 |

### SCYN segmentation is fast

Recall that efficient processing of scRNA-seq data is essential, especially in today’s thousands of single cells throughput. To evaluate the efficiency of SCYN against SCOPE, we measured the segmentation task CPU running time of SCYN and SCOPE on T10, T16M, T16P, 10x 10% spike-in, 10x 1% spike-in, and several simulation data sets (90-1, 90-2, 2000-1, 2000-2, 2000-3, 2000-4, and 2000-5), with the cell number ranging from 48 to around 2000. We respectively ran SCYN and SCOPE on each dataset ten times and calculated the mean CPU running time. As illustrated in Table 3 and Figure 6, the CPU consuming time of SCYN is almost linear in log scale with the increase of cell number. However, the CPU time of SCOPE rises dramatically when the cell number goes to hundreds or thousands. For instance, for large datasets with 2k cells, SCYN is around 150 times faster than SCOPE, SCYN finished the tasks within eight minutes, while SCOPE is unable to scale 2k cells within 16 hours. In all, SCYN is super fast in respective of datasets scale up to hundreds or thousands.

**Table 3.** benchmark for runtimes (Minutes)

| Sample | Cell Number | SCYN | SCOPE | Fold change on time |
| --- | --- | --- | --- | --- |
| T10 | 99 | 2.917 | 46.995 | 16.111 |
| T16M | 48 | 2.566 | 21.94 | 8.55 |
| T16P | 52 | 2.786 | 23.927 | 8.588 |
| 90-1 | 93 | 2.73 | 44.14 | 16.168 |
| 90-2 | 92 | 2.769 | 40.415 | 14.596 |
| 10X-1% spike-in | 1056 | 3.598 | 485.768 | 135.011 |
| 10X-10% spike-in | 462 | 2.615 | 208.854 | 79.868 |
| 2000-1 | 2173 | 6.714 | 1147.658 | 170.935 |
| 2000-2 | 2214 | 7.602 | 1122.881 | 147.709 |
| 2000-3 | 1722 | 6.817 | 947.66 | 139.014 |
| 2000-4 | 1909 | 8.139 | 1122.335 | 137.896 |
| 2000-5 | 2048 | 7.128 | 1118.038 | 156.852 |

**Fig. 6.**
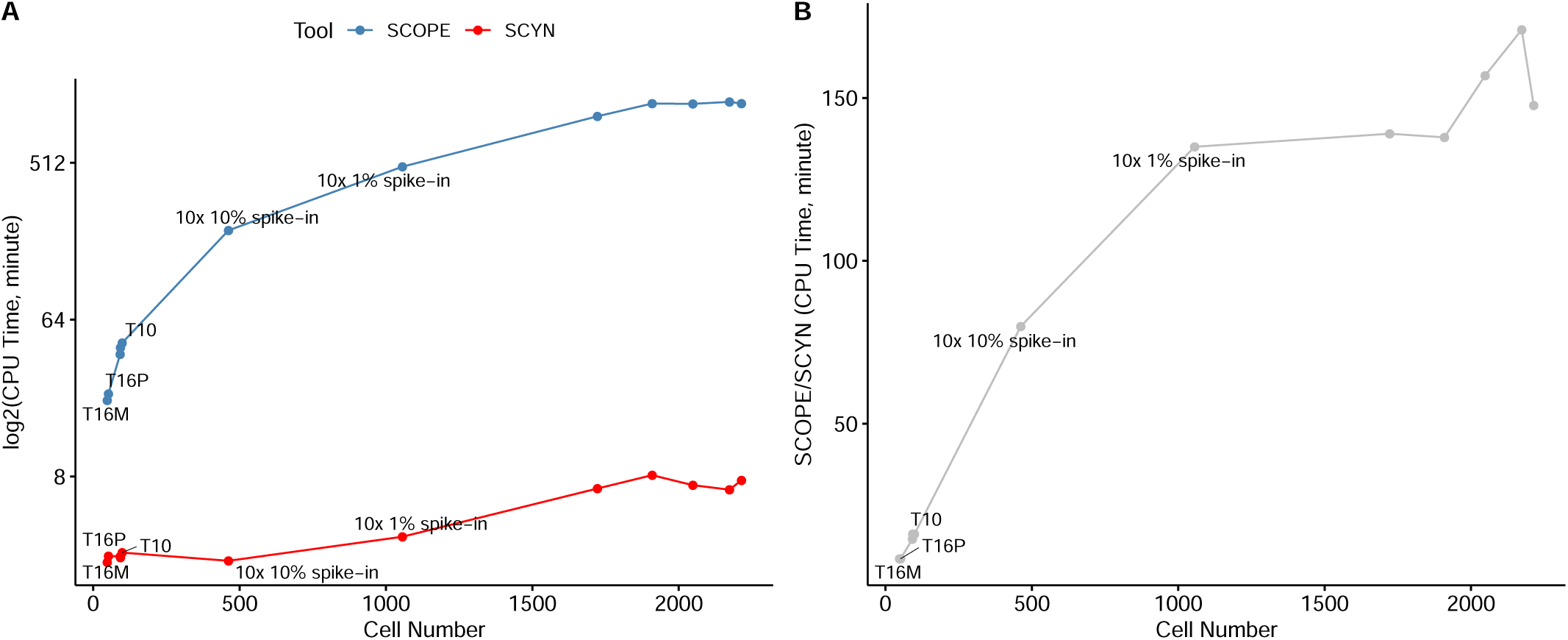
Runtime performance of SCYN. (A) CPU time of SCYN and SCOPE on different cell number scale, respectively. (B) CPU time fold change of SCOPE against SCYN on different cell number scale.

### SCYN segmentation has better mBIC values

SCYN is fast because we only adopt the simplified version (see Equation 1 in Method) of total SCOPE-mBIC (31) as the objective of segmentation and optimize it utilizing dynamic programming. Experiments on synthetic datasets and real cancer datasets successfully validated the tumor intra-heterogeneity exposure efficacy of SCYN against SCOPE. Here we further evaluate SCYN optimization effectiveness against SCOPE in respective of the original SCOPE-mBIC objective. We compared SCOPE-mBIC value by adopting the segmentation results of SCYN and SCOPE on real cancer datasets T10, T16P, T16M, and 10x spike-ins. As illustrated in Figure 7A and Supplementary Figure S5A, the mBICs yielded from SCYN on samples across all chromosomes are always more massive than the mBICs produced by SCOPE, except chromosome 16 of 1% spike-in. Clearly, SCYN achieves better segmentation concerning the tedious SCOPE objective. Furthermore, as illustrated in Figure 7B and Supplementary Figure S5B, the proportions of the simplified mBIC against overall SCOPE-mBICs are overwhelming across all chromosomes, indicating all residual terms actually can be neglected without loss of accuracy. SCYN produced smaller mBIC values than SCOPE on chromosome 16 for 1% spike-in dataset, suggesting that the residual terms take effect on circumstances such as the tiny proportion of cancer cells. However, we believe that the 1% spike case is rare in scDNA sequencing samples and is invalid for downstream analysis, and the minor fluctuations of mBIC will not affect the ability of SCYN to detect subclones, as proved in the previous section.

**Fig. 7.**
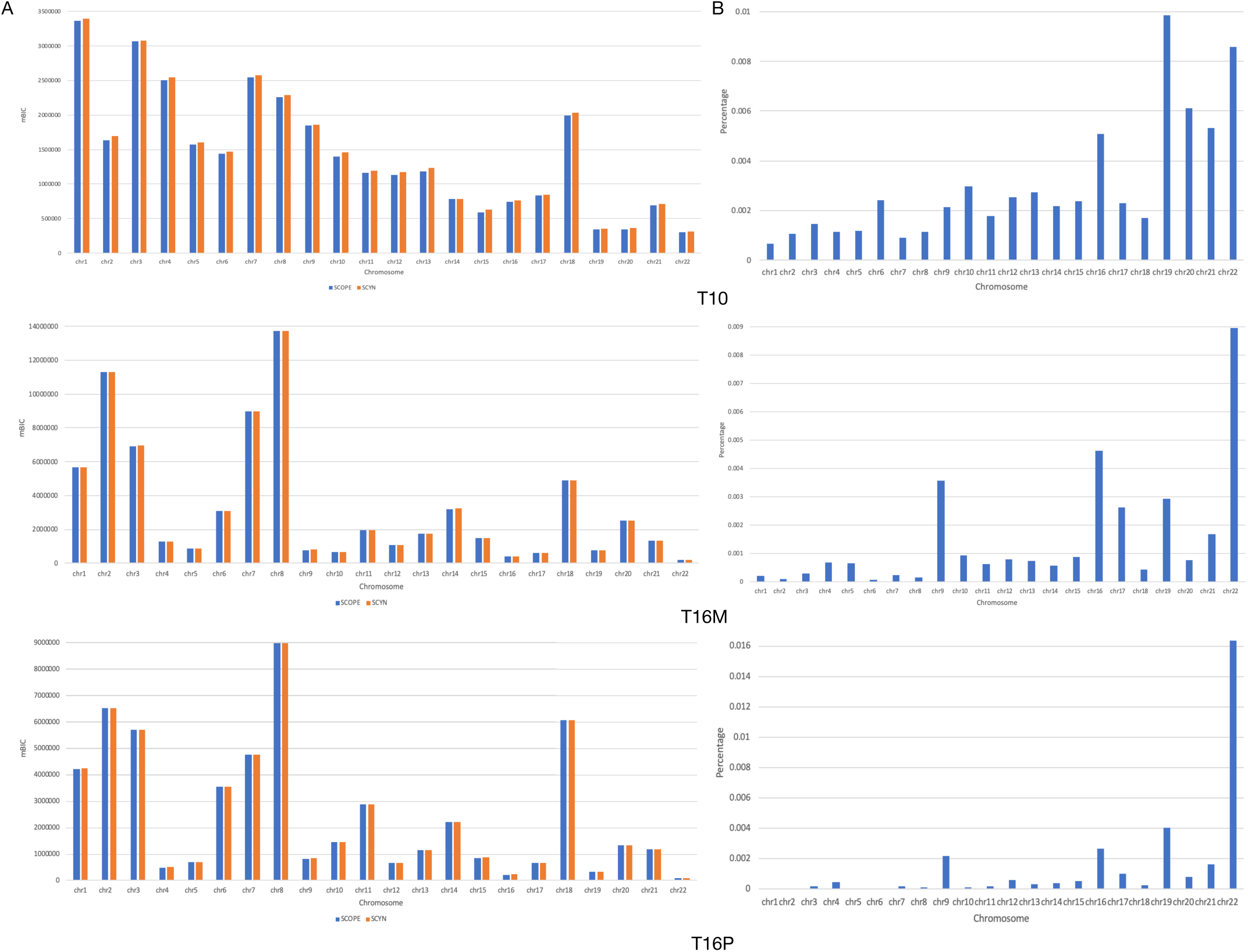
(A) SCOPE-mBIC of T10, T16M and T16P across all chromosomes generated by SCYN and SCOPE, respectively. (B) The proportion of residual terms over mBIC across all chromosomes on T10, T16M, and T16P, respectively

## Discussion

In this study, we proposed SCYN, a fast and accurate dynamic programming approach for CNV segmentation and checkpoint detection customized for single cell DNA sequencing data. We demonstrated SCYN guaranteed to resolve the precise turning points on two *in silico* datasets against SCOPE. Then we proved SCYN manifested a more accurate copy number inferring on triple-negative breast cancer scDNA data, with array CGH results of purified bulk samples as ground truth validation. Furthermore, we benchmarked SCYN against SCOPE on 10x Genomics CNV solution datasets. SCYN successfully recognizes gastric cancer cell spike-ins from diploid cells. Last but not least, SCYN is about 150 times faster than state of the art tool when dealing with thousands of cells. In conclusion, SCYN robustly and efficiently detects turning points and infers copy number profiles on single cell DNA sequencing data. It serves to reveal the tumor intra-heterogeneity.

The implementation of SCYN is wrapped in python packages https://github.com/xikanfeng2/SCYN. It provides the segmented CNV profiles and cell meta-information available for downstream analysis, such as hierarchical clustering and phylogeny reconstruction. Last but not least, the CNV profiles obtained from SCYN can be directly visualized in https://sc.deepomics.org/, which supports real-time interaction and literature-style figure downloading.

We neglected one crucial issue. Cancer scDNA-seq intensities should be regarded as a mixture of subclone cell signals with confounding of sparsity, GC bias, and amplification bias (31). The perfect CNV segmentation heavily relies on the cross-cell normalization of intensities in the first place. While we brutely adopt the normalization schema from SCOPE; there lacks a comprehensive evaluation of scDNA intensities normalization. Speaking to further work, inferring CNV profiles from single-cell RNA sequencing (scRNA-seq) is trending (11–13, 37). Incorporating DNA and RNA to profile single cell CNV segmentation might lead to tumor intra-heterogeneity to a higher resolution.

## Methods

### Data sets

#### Synthetic data

Two synthetic datasets were generated to evaluate the segmentation power of SCYN. The dimension of each dataset is 400 bins and two cells. The ground truth segmentation is (1, …, 49), (50, …, 99) (100,…, 149), (150,…, 200) for both of datasets. For the first dataset, the reads count of two cells for the four segments was designed to around (100, 100), (400, 400), (100, 100) and (400, 400), respectively. For the second dataset, the reads count of two cells for the four segments was designed to around (100, 400), (400, 100), (100, 400) and (400, 100), respectively. Random noise was applied to these reads counts.

#### Single-end Real scDNA-seq data

Two single-end breast cancer scDNA-seq datasets were downloaded from NCBI Sequence Read Archive with the SRA number of SRA018951. The raw fastq files were aligned using BWA-mem (38) to the human hg19 reference genome, and the BAM files were sorted using SAMtools (39). Picard toolkit (40) was used to remove duplicate reads. The clean BAM files were fed as the input of SCYN package.

#### Ten-X (10x) data

The 10x spike-in scDNA-seq data was collected from the 10x Genomics official dataset with the accession link https://support.10xgenomics.com/ single-cell-dna/datasets. The cell-mixed BAM files were demultiplexed to cellular BAMs according to cellular barcodes using Python scripts.

### Notations

To profile the CNV along genomes, first, we partition the genome into fix-size bins. Assume the number of bins as *m*. If the number of cells is *n*, then the input matrices, *Y*_*m*×*n*_ and *Ŷ*_*m*×*n*_, contain the raw and normalized reads counts, respectively; that is, *Y*_*i,j*_ includes the number of raw reads count belong to bin *i* at cell *j* and *Ŷ*_*m*×*n*_ contains the number of normalized reads count belong to bin *i* at cell *j*, where 1 ≤ *i* ≤ *m* and 1 ≤ *j* ≤ *n*.

### Segmentation

The first task is to partitioning the bins into segments to optimize an objective function. Here, we choose the objective function to maximize the simplified version of modified Bayesian information criteria (mBIC) proposed by Wang *et al*. (31).

To calculate the simplified mBIC, we need to partition the sequence of bins into 𝓁 segments *s*_1_, *…, s*_*𝓁*_, where *s*_*k*_ = (*i*_*k-*1_ + 1, *…, i*_*k*_), *k*_0_ = 0 ≤ *k*_1_ *< k*_2_ *< … < k*_*𝓁*_ = *n*. Denote the number of bins in segment *s*_*k*_ as |*s*_*k*_| With the partitioning, we can calculate two matrices 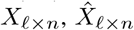, where 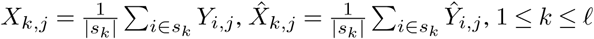. Given a segmentation *S* = (*s*_1_, *…, s*_*𝓁*_), its simplified mBIC is calculated as

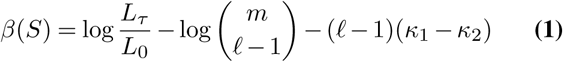

where log 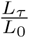 is the generalized log-likelihood ratio, *κ*_1_ and *κ*_2_ are two pre-defined constants and

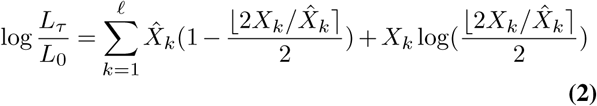

For more details on the interpretation of the terms in mBIC, we refer the readers to Wang *et al*. (31). Our objective here is to find a segmentation *S*_*opt*_ such that *β*(*S*_*opt*_) is maximized.

### Optimal algorithm

Let *β*(*k, i*) store the simplified mBIC value for the optimal segmentation which partitions bins 1, *…, i* into *k* segments. Associated with *β*(*k, i*), we also store the corresponding generalized log-likelihood ratio *L*(*k, i*), which is the first term in **Equation 1**, the log-likelihood ratio *l*(*i, j*) for a single segment starting at the *i*-th bin and ending at the *j*-th bin, and the (*k –* 1)-th optimal turning point position *T* (*k –* 1, *i*) to partition bins 1, *…, i* into *k* segments. The *β*(*k, i*) is calculated by the following recursive formulations:

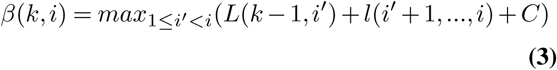

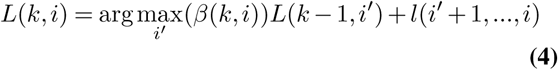

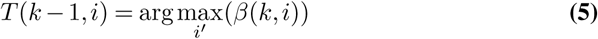

where *C* is the sum of last two terms in **Equation 1**.

As demonstrated in **Equation 3**, the value of each cell *β*(*k, i*) in table *β* can be computed based on the earlier store data *L*(*k –* 1, *i*′) and *l*(*i*′ + 1, *…, i*). The computed *β*(*k, i*) is then used to incrementally with *k* and *i* to compute the correct values of *β*. Clearly, the values of *β* and *L* for one segment can be initialized to equal to *l*.

The values of *β* can be stored in a two dimensional array, i.e., a table. The procedure for computing the table *β* is also displayed in **Algorithm 1**. The table *β* will be constructed starting from a single segment *β*(1, *i*), and moving towards more segments *β*(*k, i*). The *β*(1, *i*) and *L*(1, *i*) are initialized to *l*(1, *i*) and *T* (0, *i*) is initialized to 0 when there is only one segment. When computing a cell *β*(*k, i*)(*k >* 1), we will checks all possible *i*′, (*k* ≤ *i*′ *< i*) and compute all values of (*L*(*k* − 1, *i*′) + *l*(*i*′ + 1, *…, i*) + *C*) and *β*(*k, i*) is determined by max_(*L*(*k*−1,*i*′)+*l*(*i*′+1,…,*i*))+*C*_. Processing the bins form in increasing order on length guarantees that the final optimal segmentation can be detected when *i* is equal to the total number of bins *m*. At the last, the positions of *k –* 1 turning points are stored in table *T*.

### Backtracking

The backtracking process of finding the positions of the optimal turning points is demonstrated in **Figure 1**B. Let the table at the left-side of **Figure 1B** as *T*, where *i* and *j* are the indexes of turning points and bins respectively. *T* (*i, j*) is the position of the *i*-th optimal turning point for a segment *s*(0, *j*). The optimal total turning points number is determined by the maximum value of *β*(*i, m*), where m is the total number of bins. Then the positions of the optimal turning points can be found by the following formulation:

#### Algorithm 1 Computing the table *β*

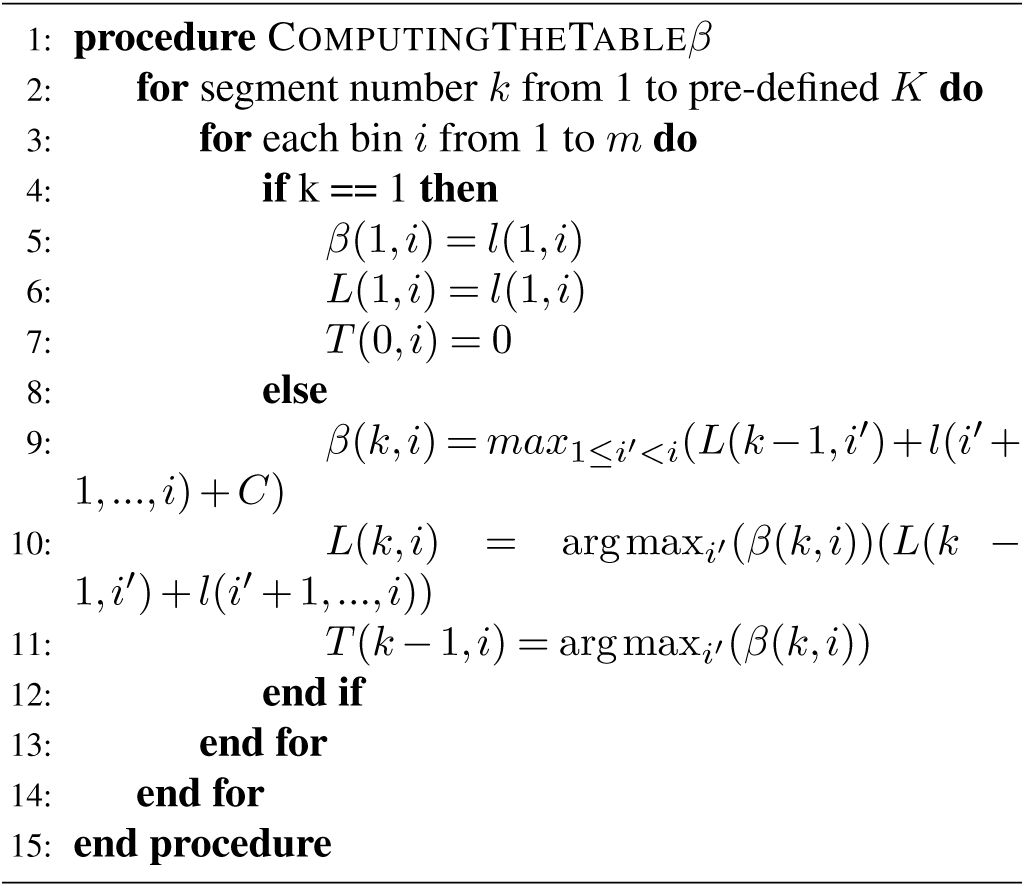

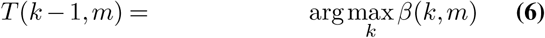

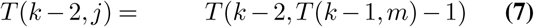

where k is the total segmentation number (1 *< k* ≤ *K*), j is the index of bin and m is the total number of bins.

### Time complexity

The time complexity of this algorithm is *O*(*m*^2^*n* + *m*^2^*k*), where m is the total bin number, n is the total cell number and k is the total segment number. The time complexity of calculating each *l*(*i, j*) is *O*(*n*) and we need to go over *O*(*m*^2^) possible segments for *m* bins. Therefore we need to *O*(*m*^2^*n*) time to construct the table *l*. For a given segments number *k*, we need to calculate *O*(*m*) possible (*L*(*k*1, *i*′) + *l*(*i*′ + 1, *…, i*)) values to get the maximum *L*(*k, i*) for *m* possible *i*, total *O*(*m*^2^) times. The time complexity for calculating the table *L* is *O*(*m*^2^*k*). In conclusion, the time complexity of our algorithm is *O*(*m*^2^*n* + *m*^2^*k*).

### Benchmark settings

SCOPE is a state-of-the-art tool for single cell CNV calling. We followed the steps in SCOPE README tutorial to perform the call CNV tasks in all datasets and the default parameters were used in all experiments. For SCYN, the function ‘call()’ was used and all parameters were set to default values. For running time analysis experiments, all experiments were run on a Dell server with an Intel(R) Xeon(R) CPU E5-2630 v3 with a clock speed of 2.40GHz. The mean value of 5 independent runs was regarded as the final running time for each tool.

## Availability of data and materials

The data and source code included in this study can be found in https://github.com/xikanfeng2/SCYN. The visualisation tools are hosted on https://sc.deepomics.org/.

## List of abbreviations

CNV: Copy Number Variation
scDNA-seq: Single Cell DNA sequencing
scRNA-seq: Single Cell RNA sequencing
aCGH: array Comparative Genomic Hybridization
CBS: Circular Binary Segmentation
HMM: Hidden Markov Model
ARI: Adjusted Rand Index
NMI: Normalized Mutual Information
JI: Jaccard Index
mBIC: modified Bayesian information criteria

## Ethics approval and consent to participate

Not applicable.

## Consent for publication

Not applicable.

## Acknowledgements

Not applicable.

## Author’s contributions

SCL. conceived the idea and supervised the project.

XF, LC, SCL discussed the algorithm and designed the experiments.

XF implemented the code and conducted the analysis.

YQ, RL, CL visualized the CNV profiles.

LC, XF drafted the manuscript.

SCL revised the manuscript.

All authors read and approved the final manuscript.

