## Supplementary File for "SCYN: Single cell CNV profiling method using dynamic programming"

March 27, 2020

#### Contents

**1 Supplementary Figures** **2**

### 1 Supplementary Figures

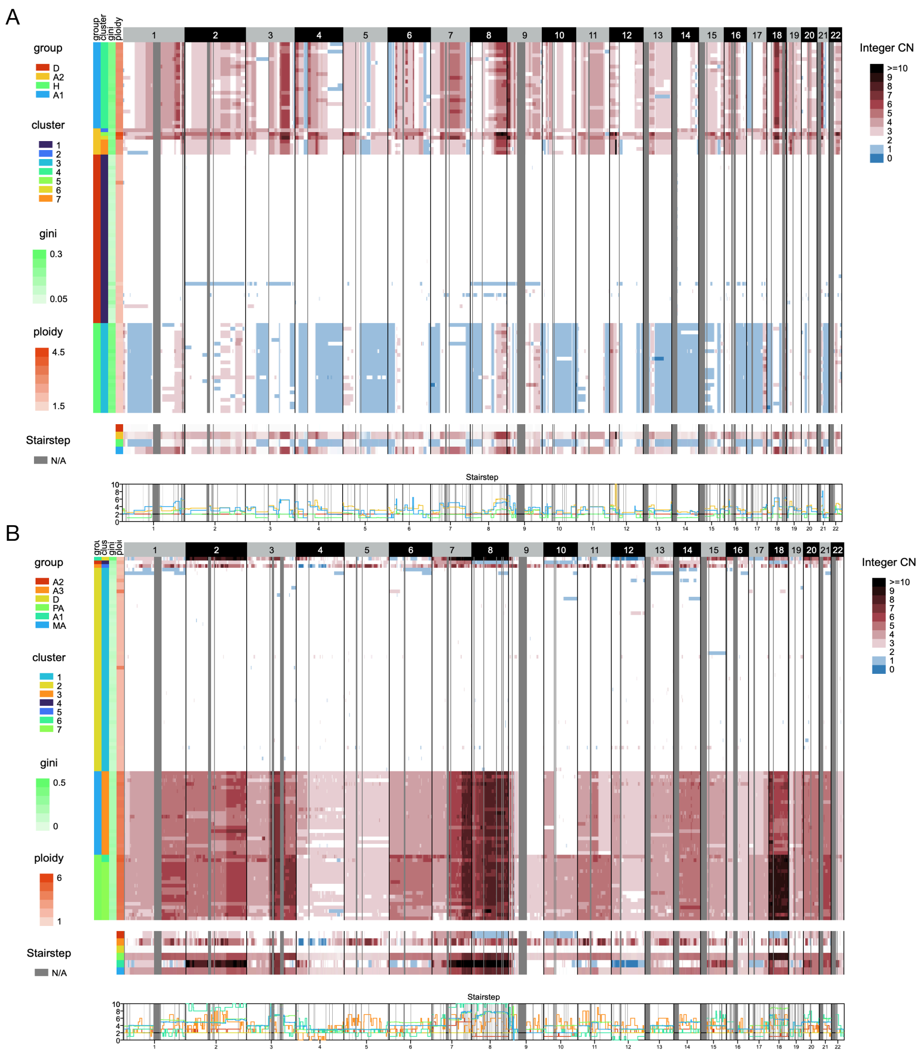


Figure S1: (A-B) Whole genome CNV profiles called by SCOPE on T10 and T16, respectively.

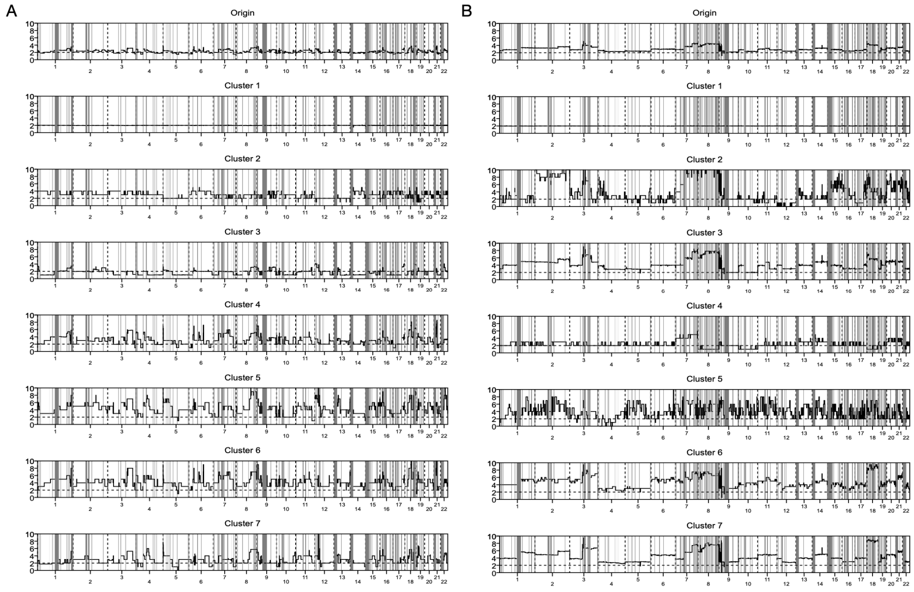


Figure S2: (A-B)SCYN ploidy stairstep plot of T10 and T16, respectively.

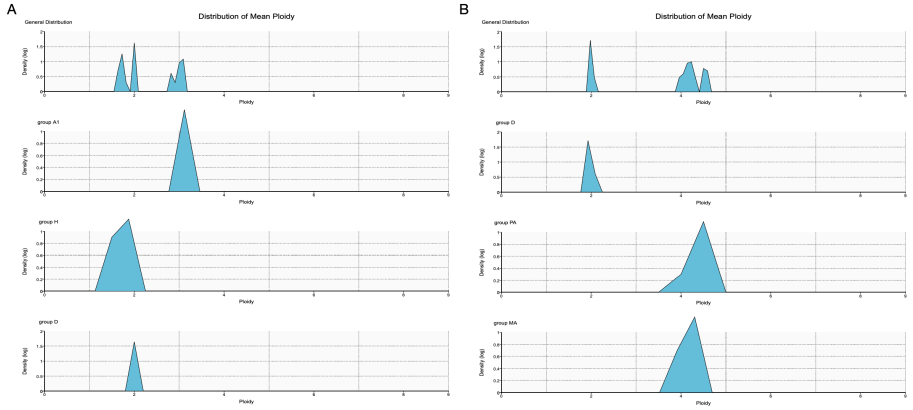


Figure S3: (A-B)SCYN mean ploidy distribution of T10 and T16, respectively.

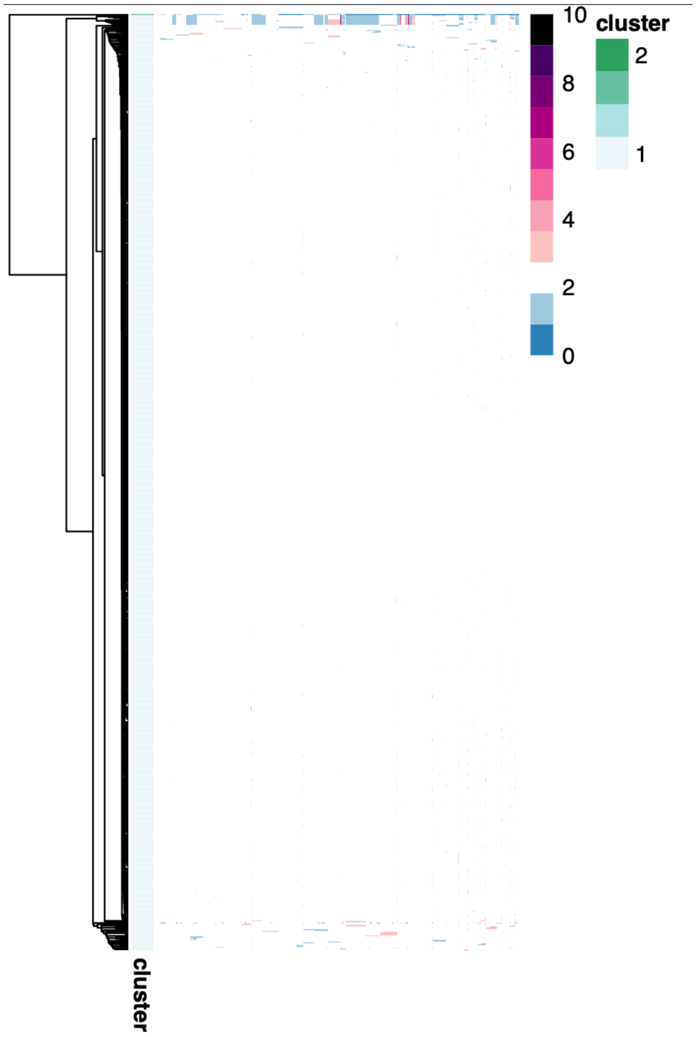


Figure S4: Whole genome CNV profiles called by SCYN on 10x 1% spike-in.

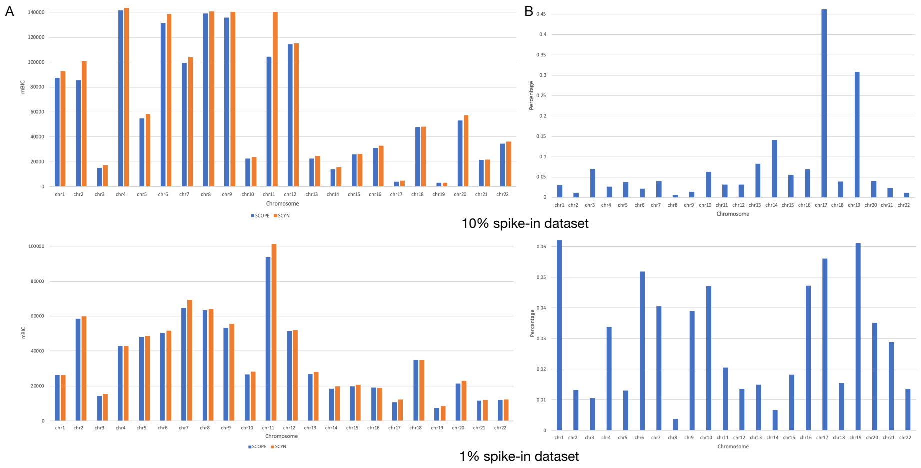


Figure S5: (A) SCOPE-mBIC of 10% and 1% spike-ins across all chromosomes generated by SCYN and SCOPE, respectively. (B) The proportion of residual terms over mBIC across all chromosomes on 10% and 1% spike-ins, respectively
